# Microbiomes and host genetics provide evidence for ecological diversification among Caribbean members of the sponge genus *Ircinia* Nardo, 1833

**DOI:** 10.1101/2020.09.04.282673

**Authors:** Joseph B. Kelly, Robert W. Thacker

## Abstract

Sponges live in symbioses with microbes that allow the hosts to exploit otherwise inaccessible resources. Given the potential of microbiomes to unlock new niche axes for the hosts, microbiomes may facilitate evolutionary innovation in the ecology of sponges. However, the hypothesis that ecological diversification evolves via the microbiome among multiple, closely related sponge species living in sympatry is yet untested. Here, we provide the first test of this hypothesis within *Ircinia*, a genus possessing diverse and abundant microbiomes that engage their hosts in nutritional symbioses. We used genome-wide SNP data (2bRAD) to delimit genetic species boundaries using BFD* among four *Ircinia* growth forms that putatively constitute distinct species and two nominal species, *I. campana* and *I. strobilina*. We also evaluated the performance of two single-locus genetic barcodes, CO1 and ITS, in resolving *Ircinia* species boundaries. We then used 16S rRNA metabarcoding to test whether the genetic species units uncovered by BFD* harbor microbiomes that are compositionally unique within each host lineage and distinct relative to seawater microbial communities. BFD* recovered genetic species boundaries that are generally reflected in the morphological differences of the growth forms and upheld the species designations of *I. campana* and *I. strobilina*, whereas CO1 and ITS provided comparatively little species-level phylogenetic resolution. The microbiomes were found to be compositionally distinct relative to seawater microbial communities, conserved within host lineages, and non-overlapping relative to the microbiomes of other host lineages. These results support a model by which microbiomes underly ecological divergence in resource use among closely related sponge species. This research provides insights into the roles of microbiomes in ecological speciation of sponges and sets the groundwork for further investigation of adaptive radiations in sponges.

## Introduction

Microbiomes, the assemblages of symbiotic microbes that live in close association with host organisms, are universal features of eukaryotes. Given the ubiquity of their presence and their impacts on fundamental biological processes of their hosts, microbiomes have experienced a growing appreciation in the scientific community over the past few decades. Their influences can be seen across multicellular organisms and include, for example, the development of immune systems in humans (Belkaid & Hand, 2014), predator evasion via bioluminescence in the Hawaiian bobtail squid (*Euprymna scolopes* Berry 1913) (Jones & Nishiguchi, 2004), and the mediation of body contractions in *Hydra* Linnaeus, 1758 (Murillo-Rincon et al., 2017). Among the roles of microbiomes is one that appears to be particularly broad in terms of the number and span of eukaryotic host clades for which it has been documented: nutrition (Akman Gündüz & Douglas, 2009; Graf, 1999; Semova et al., 2012; C. Wilkinson & Cheshire, 1990; Yuen et al., 2019; Zimmer & Bartholmé, 2003). The specific mechanisms by which microbes enable the provision of nutrients for their hosts differs among symbioses and is largely contingent on the trophic strategies of the microbes; fermentative gut prokaryotes of ruminants break down plant material that is undigestible by the host (Yeoman & White, 2014), photoautotrophic dinoflagellates use light and host-derived CO_2_ to produce sugars for zooxanthellate hermatypic corals (Davy et al., 2012); and rhizosphere bacteria perform nitrogen fixation to convert atmospheric nitrogen (N_2_) to ammonia (NH_3_), a biologically available nitrogen source for legumes (Kuypers et al., 2018).

Despite the biochemical diversity of nutrient exchanges from microbiomes, a commonality can be found among many of them in that they unlock nutrient-based resource axes that are difficult or impossible for the host organism to traverse alone: the digestive enzymes of mammals are unable to break down complex carbohydrates of roughage, metazoans lack the ability to photosynthesize, and plants cannot fix N_2_ to create NH_3_. By enabling their hosts to explore new trophic space, microbiomes may facilitate ecological diversification. The extent to which microbial symbioses can drive the evolution of ecological diversity is perhaps best exemplified by the insect order Hemiptera Linnaeus, 1758, in which microbiomes appear to have aided hosts in shifting to new food species by producing nutrients that are either lacking from the food source or cannot be provisioned by the insect’s own metabolism (Sudakaran et al., 2017). The evolutionary lability of microbiome-mediated transitions in resource use of hemipterans has resulted in multiple adaptive radiations of host plant specialists, outlining the role of microbiomes as key innovations (Heard & Hauser, 1995; Janson et al., 2008). Given that microbiomes can exert strong fitness influences on host clades across the eukaryotic tree of life and in numerous biological contexts, the role of microbiomes as conduits for ecological speciation (Brucker & Bordenstein, 2012) might be a common evolutionary force driving diversification.

Host-microbiome symbioses abound in the aquatic environment and form the foundations of many hosts’ biologies. Such is the case in sponges (phylum Porifera Grant, 1836), one of the oldest extant metazoan phyla, which first acquired symbiotic associations with microbes early in their evolutionary history (∼540 mya) (Brunton & Dixon, 1994). The sponge-microbe symbiosis has had profound impacts on the ecology of sponges due to the microbes’ roles in the exploitation of resources that are otherwise inaccessible to the hosts. The metabolisms of sponge-dwelling microbes that hold the potential to supplement their hosts’ energetic budgets encompass several microbial trophic metabolisms such as photosynthesis (Erwin & Thacker, 2007; C. J. Freeman & Thacker, 2011), the fermentation of organic compounds (Hentschel et al., 2006), sulfur oxidation (Jensen et al., 2017), methane oxidation (Rubin-Blum et al., 2019), and nitrogen fixation (C. R. Wilkinson & Fay, 1979). These processes have been demonstrated experimentally to have a positive effect on host sponge fitness, as is the case with photosynthesis (C. J. Freeman & Thacker, 2011), have strong correlative evidence for supplementing host nutrition, as has been observed in methanotrophic symbionts (Rubin-Blum et al., 2019), or likely affect sponge fitness positively based on the physiological exchanges documented in other host clades that possess symbionts performing similar biochemical functions. These include the sulfur oxidizing symbionts of lucinid clams (Yuen et al., 2019) and vestimentiferan tube worms (Stewart & Cavanaugh, 2006), and the aforementioned examples of the nitrogen fixation in legumes (Kuypers et al., 2018) and fermentation in ruminants (Yeoman & White, 2014).

Many sponge species possess microbiomes with taxonomic compositions that are conserved within host species and are distinct relative to the microbiome compositions of other sponge species (Thomas et al., 2016). Isotopic data represented by sympatric, though phylogenetically distant host taxa support the hypothesis that this pattern of beta diversity among the microbiome compositions translates to divergent niche inhabitation among sponge species and potentially facilitates their coexistence on the same reef (C. J. Freeman et al., 2020). However, tests of ecological divergence among congeneric or incipient sponge species are few in number because the datatypes required to provide the necessary level of phylogenetic resolution among the host lineages, such as genome-wide SNP data, are seldom used in conjunction with metabarcoding or metagenomic censuses of microbiomes in sponges. To our knowledge only two studies have implemented such a study design by evaluating both microsatellite and 16S rRNA data, one in *Ircinia campana* Lamarck, 1814 (Griffiths et al., 2019) and the other in *Cliona delitrix* Pang, 1973 (Easson et al., 2020). In both systems, a positive correlation between genetic distance and beta diversity in microbiome composition was observed, suggesting that the development of distinct microbiome compositions coincides with genetic divergence between sponge lineages prior to or contemporaneously with the onset of speciation. However, the hypothesis that diversification in resource use is driven by microbiomes in multiple closely related sponge lineages inhabiting the same geographic area remains unexplored.

A suitable system to investigate this question exists in Caribbean *Ircinia* Nardo 1833. Sponges in this genus host symbiotic prokaryotic communities that can be four orders of magnitude more abundant than and compositionally distinct relative to prokaryotic communities in the surrounding seawater, reflecting a pervasive role of microbiomes in *Ircinia* physiologies (Erwin et al., 2012; Hardoim & Costa, 2014). Additionally, members of this genus rank among the most productive in shallow-water Caribbean benthic habitats due to the substantial rates of photosynthesis by their microbiomes (C. Wilkinson & Cheshire, 1990; Wulff, 1994). Of the four currently described shallow water Caribbean *Ircinia* at least two, *I. campana* and *I. felix* Duchassaing & Michelotti, 1864, are likely mixotrophs, gaining energy from symbiont photosynthesis as well as through heterotrophic filter feeding (Erwin & Thacker, 2007; C. Wilkinson & Cheshire, 1990). Previous work on microbial metabolisms in *I. felix* suggests that bacterial photosynthesis and nitrogen metabolism play important roles in subsidizing host nutrition (Weisz et al., 2007). *I. felix* microbiomes may also perform phosphorus transformations, although the link to host nutrition has not yet been made (Archer et al., 2017). The microbes of Caribbean *Ircinia* are also involved in the generation of quorum sensing signals and inhibitors that may prevent overgrowth by pathogens (Mohamed et al., 2008; Quintana et al., 2015). Given the importance of symbiotic microbes in *Ircinia*, investigating host-microbe associations within this genus could help illuminate the roles that microbiomes play in generating ecological diversity among host lineages of aquatic organisms.

Several studies have reported additional putative Caribbean *Ircinia* species that are recognizable by differences in macroscopic features of their bodies’ growth morphology (Diaz, 2005; Erwin & Thacker, 2007; Rützler et al., 2000; van Soest, 1978), suggesting that the recognition of only four species is an underrepresentation of the taxonomic richness of this genus in the Caribbean. These growth forms appear to occupy distinct ecological roles, evidenced, for example, by differences in chlorophyll-*a* concentrations in their mesohyl (Erwin & Thacker, 2007). However, it is unknown if the growth forms are distinct species and, if they are separate species, what biological factors contribute to their diversification. Given the likely influences of microbiomes on *Ircinia* fitness, it is reasonable to postulate that host traits promoting the stability of an advantageous symbiotic microbial community will be evolutionarily favored. Additionally, if microbes enable sponges to exploit different resources, potentially facilitating local adaptation and alleviating resource competition, then the relative abundances of physiologically impactful microbial groups will differ among host species.

In this study, we investigated whether microbiomes contribute to ecological divergence among closely related *Ircinia* lineages inhabiting the same geographic locale, a stretch of shallow water coastal environment in Bocas del Toro, Panama. We levied genome-wide SNP data to delimit species boundaries and to identify patterns of hybridization among four *Ircinia* growth forms and two nominal species, *I. campana* and *I. strobilina* Lamarck, 1816. We then investigated patterns of ecological divergence among the species recovered using the genome-wide SNP data by testing for differences among their microbiome compositions, which we inferred via 16S rRNA metabarcoding. Additionally, we analyzed nucleotide sequence data from two single-locus barcodes that are commonly used to perform population and species-level genetic studies in sponges, with the goal of asking whether these loci provide sufficient phylogenetic resolution and accuracy to delimit sponge species.

## Materials & Methods

### Specimen Collection

Thirty *Ircinia* specimens representing four growth and two nominal species, *I. campana* and *I. strobilina*, were collected from Bocas del Toro, Panama, during July 2016 from three sites (Table S1, Fig. 1). Three of the growth forms have globose or irregularly massive body morphologies. The first, Massive A pink, has 2 mm-high rounded conules, oscula no larger than 1.2 cm in diameter, and a pink exterior. The surface texture of the second growth form, Massive A green, resembles that of Massive A pink although the conules are shorter (1.5-1.75 mm). The size of Massive A green’s oscula are roughly the same size as those of Massive A pink; however, the exterior is green. The third growth form, Massive B, differs from the first two by possessing higher conules (3-5 mm), a tan exterior, and larger oscula that can reach 1.5 cm in diameter. The fourth growth form, referred to as Encrusting, grows as a green undulating mat and occasionally possesses digitate projections of the body. Like the massive growth forms, Encrusting also has low conules that are no larger than 2 mm. Each growth form was found in only one habitat; Massive A pink was found on the mangrove roots of Inner Solarte (9°18’20.9”N 82°10’23.5”W), Massive A green and Massive B were found in the seagrass-dominated habitat of STRI Point (9°21’06.1”N 82°15’32.4”W), and Encrusting was found on the patch reefs at Punta Caracol (9°22’37.6”N 82°18’08.3”W). Specimens of *I. campana* and *I. strobilina* were both collected alongside the growth forms at Punta Caracol and STRI Point. All habitats fall within a 8.1-km radius (Fig. 1).

**Figure 1.**
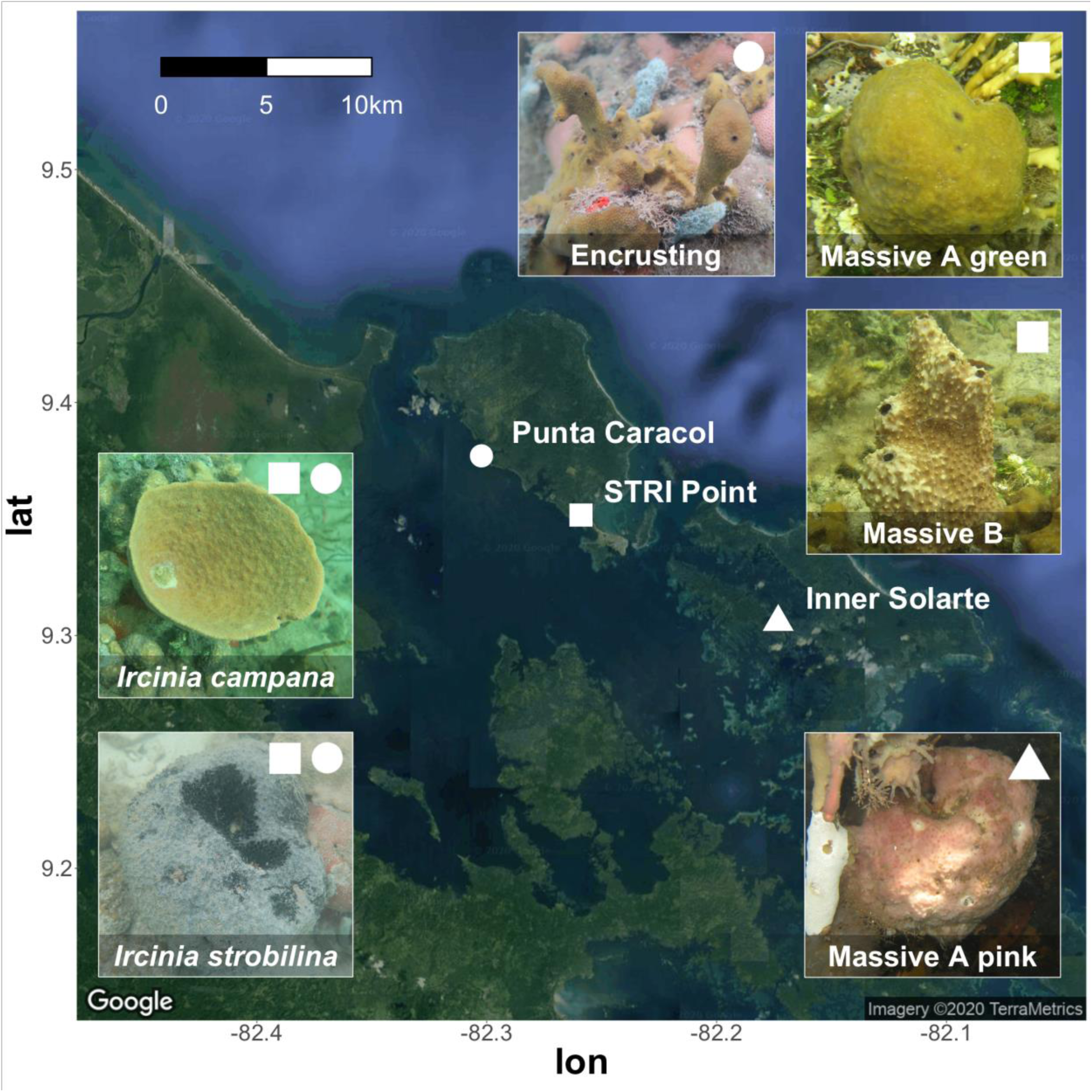
Map depicting sampling locations of Panamanian *Ircinia*. Specimens of Massive A pink were collected from the *Rhizophora* prop roots at the Inner Solarte mangrove hammock (9°18’20.9”N 82°10’23.5”W), specimens of Massive A green and Massive B were collected from the *Thalassia*-dominated seagrass habitat of STRI Point (9°21’06.1”N 82°15’32.4”W), and specimens of the Encrusting growth form were collected from the patch reef habitat Punta Caracol (9°22’37.6”N 82°18’08.3”W). Specimens of *I. campana* and *I. strobilina* were also collected from STRI Point and Punta Caracol.

Specimens of the same growth form or nominal species were collected at least 3-20 meters apart so as to avoid sampling of clones and to prevent sampling of sponges from the same microenvironment, which could result in similar influences on the microbiomes and magnify metrics of beta diversity among the growth forms’ microbiome compositions. Upon collection, thumb-sized tissue subsamples were excised from the sponges using surgical scissors and subsequently transported to the Smithsonian Tropical Research Station in Bocas del Toro for immediate processing. Samples were fixed for molecular data collection in 90% EtOH, followed by two EtOH replacements after 24 h and 48 h. Additional vouchers were made in 4% paraformaldehyde for morphological analysis; however, the measurements of internal anatomical features are reported in a separate manuscript (Kelly & Thacker 2020, *in review*). Seawater specimens (0.5 L) were collected adjacent to the sponges, transported in opaque (brown) Nalgene bottles, vacuum filtered through 0.2-μm Whatman filter papers, and stored in RNA later.

### DNA Extraction and Next-Generation Sequencing Library Preparation

DNA was extracted from the exterior (outermost 2 mm) tissue and water filters using the DNeasy PowerSoil Kit (Qiagen) and from the interior tissue (at least 2 mm away from the outermost edge of the specimen) using the Wizard Genomic DNA Purification Kit (Promega), following the manufacturer’s instructions. DNA was extracted from both the interior and exterior portions of the sponges’ bodies to optimize the type of data that was generated from each tissue region. Other researchers have reported a 2 mm-wide dark band in Caribbean *Ircinia* in the outermost region of tissue cross-sections that corresponds to cyanobacteria (C. Freeman & Gleason, 2010). We observed this band in the current *I. campana* as well as in the growth forms, and thus by extracting DNA from tissue beyond this band and towards the interior of the sponge body, we sought to produce DNA extractions that have higher ratios of concentrations of host DNA relative to microbial DNA. Conversely, by extracting DNA from the exterior of the sponge, we sought to produce DNA extractions with not only higher concentrations of microbial DNA relative to host DNA, but also to produce DNA extractions of microbes that are exposed to light, one of the primary resource axes of *Ircinia* microbiomes (Erwin & Thacker, 2007; C. Wilkinson & Cheshire, 1990), in addition to other resource axes that are relevant to chemoautotrophic microbes.

A RADseq library was constructed following the 2bRAD workflow using the Wizard Genomic DNA Purification isolations obtained from the interior portion of the sponges and the *Alf1* restriction enzyme (S. Wang et al., 2012). 2bRAD was chosen to generate host SNP data as it can target a subset of the total pool of loci in a genome using a reduced-representation library preparation scheme, which helps ensure sufficient sequencing coverage at the cost of obtaining fewer loci (S. Wang et al., 2012). By adopting such a methodology, sufficient loci can be recovered to perform a multispecies coalescent test of species boundaries via Bayes Factor Delimitation using genome-wide SNP data (BFD*), a method that allows for the evaluation of several species-grouping hypotheses and performs well with small SNP datasets and few individuals sampled per species (Leaché et al., 2014). A reduced representation library preparation scheme targeting 1/32 of *Alf1* restriction sites was performed using the primers 5ILL-RG (5’ CTA CAC GAC GCT CTT CCG ATC TRG 3’) and 3ILL-YG (5’ CAG ACG TGT GCT CTT CCG ATC TYG 3’).

The V4 region of the 16S rRNA subunit was amplified from the DNeasy PowerSoil Kit DNA extractions that were produced from the exterior portion of the sponges and the seawater samples using the primers 515f (5’ GTG YCA GCM GCC GCG GTA A 3’) and 806rB (5’ GGA CTA CNV GGG TWT CTA AT 3’) following protocols developed by the Earth Microbiome Project (http://press.igsb.anl.gov/earthmicrobiome/protocols-and-standards/16s/). PCR reactions were conducted in 50 uL volumes, with 25 uL of 2x HotStarTaq Master Mix, 1 uL of each of the primers at 10 uM concentration, 22 uL of water, and 1 uL of template DNA. Thermocycler conditions entailed an initial denaturing step of 95°C for 5 minutes followed by 35 cycles of the following progression: 94°C for 45 seconds, 50°C for 1 minute, and 72°C for 1.5 minutes; and completed with a 10-minute-long final elongation step.

The 2bRAD library was cleaned using the Wizard SV Gel and PCR Clean-up system, and the 16S libraries were cleaned using the AxyPrep Mag PCR Clean-up kit. All next-generation sequencing libraries were dual multiplexed using two 12-basepair Golay barcodes. All libraries were normalized prior to pooling following the quantification of nucleotide concentrations using a Qubit 3.0. Amplifications of 16S rRNA were split between three libraries, one of which was sequenced on an Illumina MiSeq instrument at the Cornell University Institute of Biotechnology, one at the Nova Southeastern University, and a third in the lab of Dr. Noah Palm at Yale University. The Cornell sequencing round also contained the complete 2bRAD library.

### Next-Generation Sequence Data Quality Filtering and Processing

The 2bRAD reads were processed once in trimmomatic v0.36 using the setting MAXINFO:40:0.8 to remove low-quality bases at the end of the read, and again using the HEADCROP:14 setting to remove the barcode and one of the two-basepair overhangs (Bolger et al., 2014). Removal of the second overhang was performed using the bbduk v38.26 function forcetrimright=31 [http://jgi.doe.gov/data-and-tools/bb-tools/]. Processed 2bRAD sequences were *de novo* assembled in Stacks v2.41 using the settings -m 2 -M 3 -n 2 (Catchen et al., 2013). For loci with more than one variant site, only the first SNP was included in downstream analyses. A dataset excluding SNPs that were not present in at least 75% of the samples was generated using the Stacks populations program. The cutoff of a locus’s inclusion in 75% of the samples was chosen, as this threshold has been demonstrated to reduce symbiont contamination in RADseq datasets to about 1.5% in anthozoans that are heavily populated by microbes (Titus & Daly, 2018).

16S rRNA reads were processed in trimmomatic v0.36 using the following parameters: TRAILING:30 SLIDINGWINDOW:5:30 MINLEN:150. Further QC steps, including the removal of chimeric sequences, and microbial operational taxonomic unit (herein OTU) inference were performed in mothur v1.39.5 (Schloss et al., 2009). To help ensure that the level of phylogenetic resolution was comparable between the two runs, the reads from the Nova Southeastern University run were trimmed to match the length of the Cornell reads (172 bps). Clustering of sequences into OTUs was performed using vsearch at the 98% clustering threshold (Rognes et al., 2016). Singleton and doubleton OTUs were removed from the dataset to further mitigate sequencing error. Taxonomic assignments of OTUs were made using the Silva v.132 database (Quast et al., 2013). Sequences identified by the classify.seqs command in mothur as mitochondria, chloroplasts, or eukaryotic 18S sequences were removed from the OTU relative abundance matrix prior to downstream analyses.

### Species Inference and Investigation of Hybridization Using Host SNP Data

Species boundaries within the host SNP data were identified using BFD* in Beast v.2.5.1 (Bouckaert et al., 2014; Leaché et al., 2014). The expected divergence prior θ was calculated as the average pairwise nucleotide diversity (alpha = 1, beta = 130) and the birth rate prior λ was sampled from a gamma distribution with the parameters (2, 200) (https://www.beast2.org/bfd/). Marginal likelihood estimates were made for each species-grouping hypothesis using path sampling analysis (Bouckaert et al., 2014). Each marginal likelihood analysis was run for 100,000 generations with 28 path steps and a pre-burn-in of 25,000 generations; likelihood estimates and trees were logged every 500 generations. Bayes Factors were calculated and compared following Kass and Rafftery (1995). Eighteen models were evaluated that represented different species grouping hypotheses, including models that represent the growth forms as phenotypes of *I. campana* and *I. strobilina*. Additionally, we included a model that was motivated by our hybridization analysis results (see below) termed “split massive B by STRUCTURE populations” whereby Massive B individuals were split between either the Encrusting or *I. campana* species group. Species tree estimation was performed in SNAPP v1.3.0 for the species grouping hypotheses that received the highest support from the BFD* analysis (Bryant et al., 2012). Each species tree was inferred using four MCMC chains, each with a length of 2 million generations and a burn-in of 25% (totaling 6 million generations post-burn-in). Likelihoods, theta estimates, and trees were logged every 1000 generations.

To investigate hybridization among the growth forms and nominal species, an ancestry analysis was performed in STRUCTURE v2.3.4 (Pritchard et al., 2000). *K* was set to range from 3 to 8, and 10 runs were performed for each *K* using an MCMC length of 200,000 with a burn-in of 50,000 generations. The true number of ancestral populations was predicted by calculating Δ*K* using the Evanno method implemented in Structure Harvester v0.6.94 (Earl & vonHoldt, 2011; Evanno et al., 2005).

### Microbiome Statistics

The raw abundance matrix of OTUs was transformed to relative abundance for all downstream statistics. The microbiomes of the host species supported by the BFD* analysis were tested for compositional differences using Permutational Analysis of Variance (PERMANOVA) based on Bray-Curtis dissimilarity, implemented in the adonis function of the R package vegan v2.5-3 (Oksanen et al., 2016). P-values were adjusted for pairwise PERMANOVAs using Benjamini-Hochberg corrections (Benjamini & Hochberg, 1995). The PERMANOVA assumption of homogeneity of variances was tested using PERMDISP (betadisper), and the cumulative contributions of OTUs to pairwise comparisons (Bray-Curtis dissimilarities) were calculated using SIMPER. The top 10 most influential OTUs (ranked by contribution to Bray-Curtis dissimilarity) in each pairwise comparison between host species were queried via BLASTn to a database of sequences derived from OTUs that are putatively vertically transmitted in *I. felix* (Schmitt et al., 2007). An OTU was designated a significant hit if it had 100% coverage to the database of 16S rRNA sequences from vertically transmitted symbionts, a percent identity > 98%, and an E-value < 1e-50. The vegan function metaMDS was used to construct an nMDS plot based on Bray-Curtis dissimilarity, and overlap in standard ellipse areas (SEAs) were calculated with the maxLikOverlap function in SIBER v2.4.1 (Jackson et al., 2011). The number of OTUs unique to sponges and seawater was calculated using the venn function in VennDiagram v1.6.20 (Chen, 2018).

### Single-Locus Genotyping of ITS and CO1

A DNA segment spanning the two Internal Transcribed Spacer (ITS) regions and the 5.8S rRNA subunit was amplified using the primers Por18hf (5’ GAG GAA GTA AAA GTC GTA ACA AGG) and dgPor28.63R (CTK ANT DAT ATG CTT AAR TTC AGC GGG T) in the following reaction mixture: 25 uL of 2x HotStarTaq Master Mix, 1 uL of each of the primers at 10 uM concentration, 10 uL of 5 M betaine, 4 uL of 25 mM MgCl_2_, 8 uL of water, and 1 uL of template DNA for a final volume of 50 uL (Redmond et al., 2013; Thacker et al., 2013). Thermal cycles for amplification of the ITS region followed an initial denaturing step at 95°C for 15 minutes followed by 35 cycles of the following progression: 94°C for 1 minute, 53.8°C for 1 minute, and 72°C for 1.5 minutes; and completed with a 10-minute-long final elongation step.

A segment of the mitochondrial *cytochrome oxidase c subunit 1* (CO1) was amplified using the primers cox1.IrcF (5’ GAT AAT GCG GYT CGA GTT GK 3’) and cox1.IrcR (5’ CTA CCG GAT CAA AGA AAG AA GTR T 3’) in the following reaction mixture: 25 uL of 2x HotStarTaq Master Mix, 2 uL of each of the primers at 10 uM concentration, 20 uL of water, and 1 uL of template DNA for a final volume of 50 uL. PCR amplification followed a cycle of an initial denaturing step at 95°C for 15 minutes, 35 cycles of the following progression: 94°C for 1 minute, 62°C for 1 minute, and 72°C for 1.5 minutes; and a 10-minute-long final elongation step. All ITS and CO1 reaction products were cleaned using the Wizard SV Gel and PCR Clean-up kit, normalized by hand after quantification using a Qubit 3.0 (Invitrogen), and sequenced using Sanger technology at the DNA Sequencing Facility at Stony Brook University. A table of next generation and sanger sequencing efforts can be accessed in Table S1.

### Gene Tree Inference and Analysis

Single-locus sequences (CO1 and ITS) were assembled into contigs using CodonCode v7.1.2 and aligned in MAFFT v7 using default gap penalties and the scoring matrix setting of 1PAM/K=2 (https://mafft.cbrc.jp/alignment/server/). Ambiguous positions were represented in sequences following the IUPAC nucleotide code. Publicly available sequences representing the non-irciniid dictyoceratids *Hyrtios erectus* (GenBank accession JQ082819.1), *Spongia officinalis* (HQ830364.1), and *Dysidea arenaria* (JQ082809.1) and the irciniids *I. felix* (JX306086 and JX306085), *I. stroblina* (JX306087, JX306088, JX306089, GQ337013, and NC013662), *I. oros* (JN655186.1), *I. dendroides* (KX866768.1), *I. variabilis* (HG816011.1), *I. fasciculata* (JN655174.1), *Psammocinia bulbosa* (JQ082836.1), *P. halmiformis* (FN552814.1), *Sarcotragus spinosulus* (MK350317.1), and *S. foetidus* (KX866772.1) were included in the CO1 alignment. Publicly available sequences representing *H. erectus* (AY613970.1), *D. arenaria* (JQ045725.1), *Spongia sp*. (KX688730), and *Vaceletia* sp. (AJ633837.1) and the irciniids *I. felix f. felix* (AJ703888.1), *I. oros* (LT935654.1) were included in the ITS alignment. The assumption of base composition homogeneity was tested using the disparity index test of pattern heterogeneity as implemented in Mega 7.0 using an MCMC of 10,000 (Kumar et al., 2016; Kumar & Gadagkar, 2001). JModelTest v2.1.7 was used to determine which model of nucleotide substitution best fit each alignment (Darriba et al., 2012). Bayesian inference of gene trees was performed in MrBayes v3.2.2 using a non-clock model (four parallel runs, each with four chains) with generation lengths sufficient for the average standard deviation of split frequencies to be consistently estimated below 0.01 (Ronquist et al., 2012). Burn-in values of 25% were used in consensus tree generation for both single-locus datasets. Reciprocal monophyly was determined by visually examining the trees.

### Data Availability

Raw reads for this study are lodged under the GenBank accession numbers [NNNN]. XML files for BFD* analysis, SNP matrices in Nexus format, single-locus alignments, photos of the sponge specimens, and the OTU matrices can be obtained at Data Dryad [NNNN]. All phylogenetic trees can be accessed at TreeBASE under project [NNNN].

## Results

### Species Boundaries Within the Host SNP Data

A total of 4,835,474 clusters corresponding to 2bRAD sequences were recovered after quality controls, which assembled into 53,431 2bRAD loci with an average per-locus read depth of 20.6 +/- 5.6x (1 s.d.). After applying population and sample constraints, 266 variant sites (SNPs) remained, each sourced from an independent locus. This volume of SNPs is commensurate with the only other RADseq dataset published to date, whereby 577 SNPs were obtained from 62 individuals of *Dendrilla antarctica* after filtering out loci that were not present in at least 40% of the individuals (Leiva et al., 2019). The percent of missing data per specimen in our final SNP matrix was 17.3% +/- 14.2%. None of the specimens displayed identical genotypes, suggesting that clones are not present in the dataset.

Species delimitation by BFD* lent highest support to two models with nearly equal marginal likelihoods: the model representing all four *Ircinia* growth forms as distinct species (hereafter species model 1) and the model that was motivated by the STRUCTURE analysis that split Massive B between Encrusting and *I. campana* while designating the other three growth forms as distinct species (hereafter species model 2) (Table 1). The topologies of the species trees for these two hypotheses were congruent, save for the absence of the Massive B terminal branch in model 2 (Fig. 2). Additionally, the nodes of both trees received high support values with the exception of the node joining *I. campana* and Massive B in model 1, which was the only node to be present in less than 50% of the posterior distribution (Fig. 2A). A number of alternative tree topologies were also present in the posterior distribution of the coalescent simulation (Fig. 2).

**Table 1.**
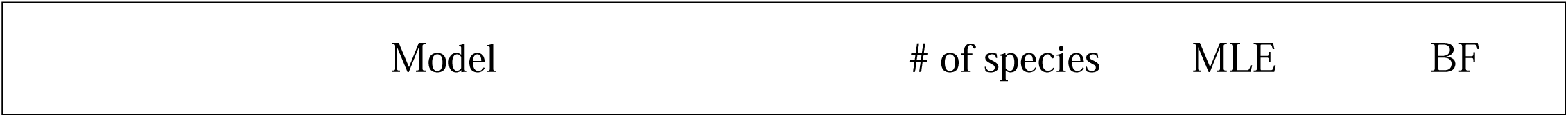
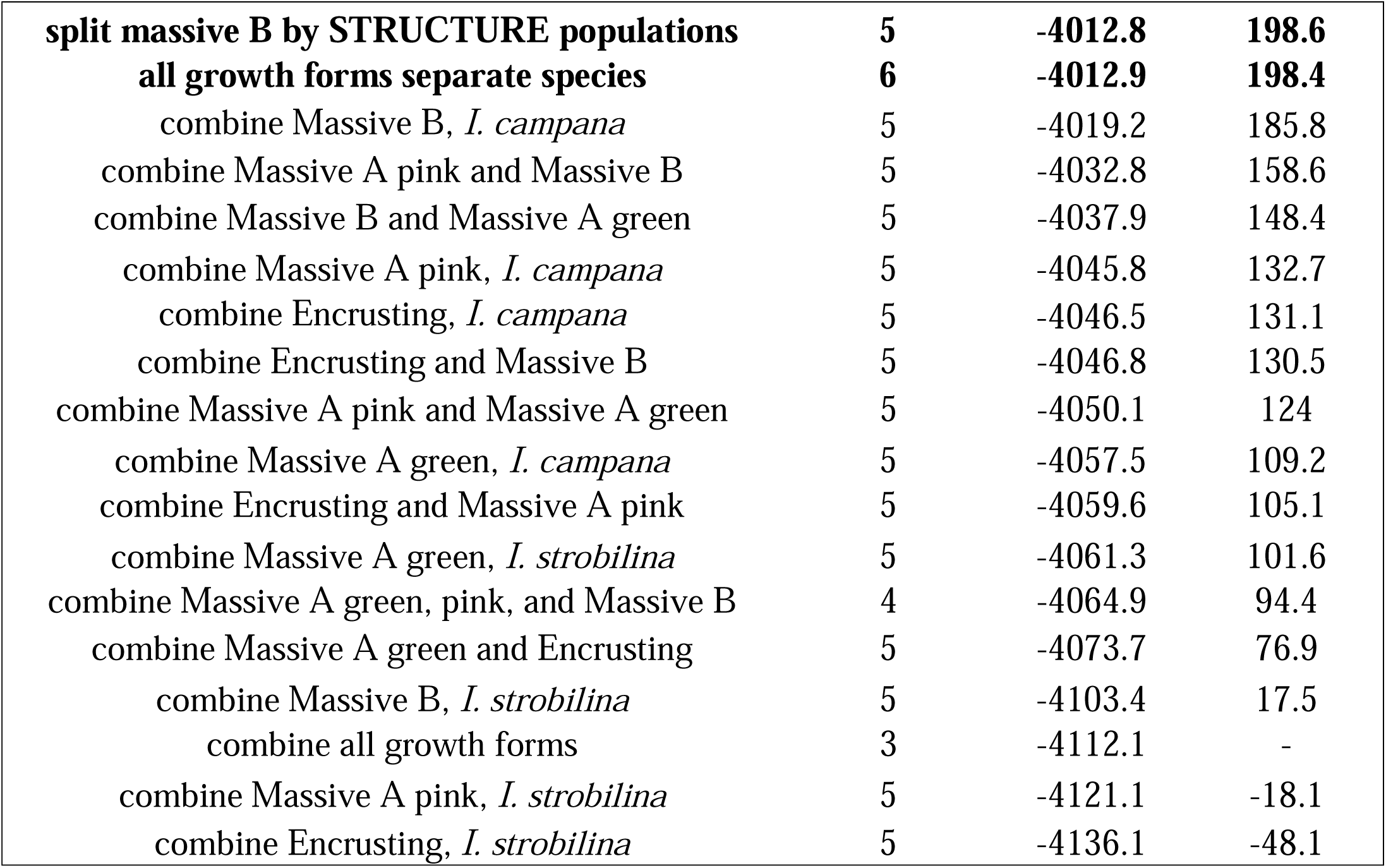
Bayes factor delimitation results. Models are listed in order of descending support with the models that received the highest support in bold font at the top.

|  |  |  |  |
| --- | --- | --- | --- |
| **split massive B by STRUCTURE populations** | **5** | **-4012.8** | **198.6** |
| **all growth forms separate species** | **6** | **-4012.9** | **198.4** |
| combine Massive B, *I. campana* | 5 | -4019.2 | 185.8 |
| combine Massive A pink and Massive B | 5 | -4032.8 | 158.6 |
| combine Massive B and Massive A green | 5 | -4037.9 | 148.4 |
| combine Massive A pink, *I. campana* | 5 | -4045.8 | 132.7 |
| combine Encrusting, *I. campana* | 5 | -4046.5 | 131.1 |
| combine Encrusting and Massive B | 5 | -4046.8 | 130.5 |
| combine Massive A pink and Massive A green | 5 | -4050.1 | 124 |
| combine Massive A green, *I. campana* | 5 | -4057.5 | 109.2 |
| combine Encrusting and Massive A pink | 5 | -4059.6 | 105.1 |
| combine Massive A green, *I. strobilina* | 5 | -4061.3 | 101.6 |
| combine Massive A green, pink, and Massive B | 4 | -4064.9 | 94.4 |
| combine Massive A green and Encrusting | 5 | -4073.7 | 76.9 |
| combine Massive B, *I. strobilina* | 5 | -4103.4 | 17.5 |
| combine all growth forms | 3 | -4112.1 | - |
| combine Massive A pink, *I. strobilina* | 5 | -4121.1 | -18.1 |
| combine Encrusting, *I. strobilina* | 5 | -4136.1 | -48.1 |

**Figure 2.**
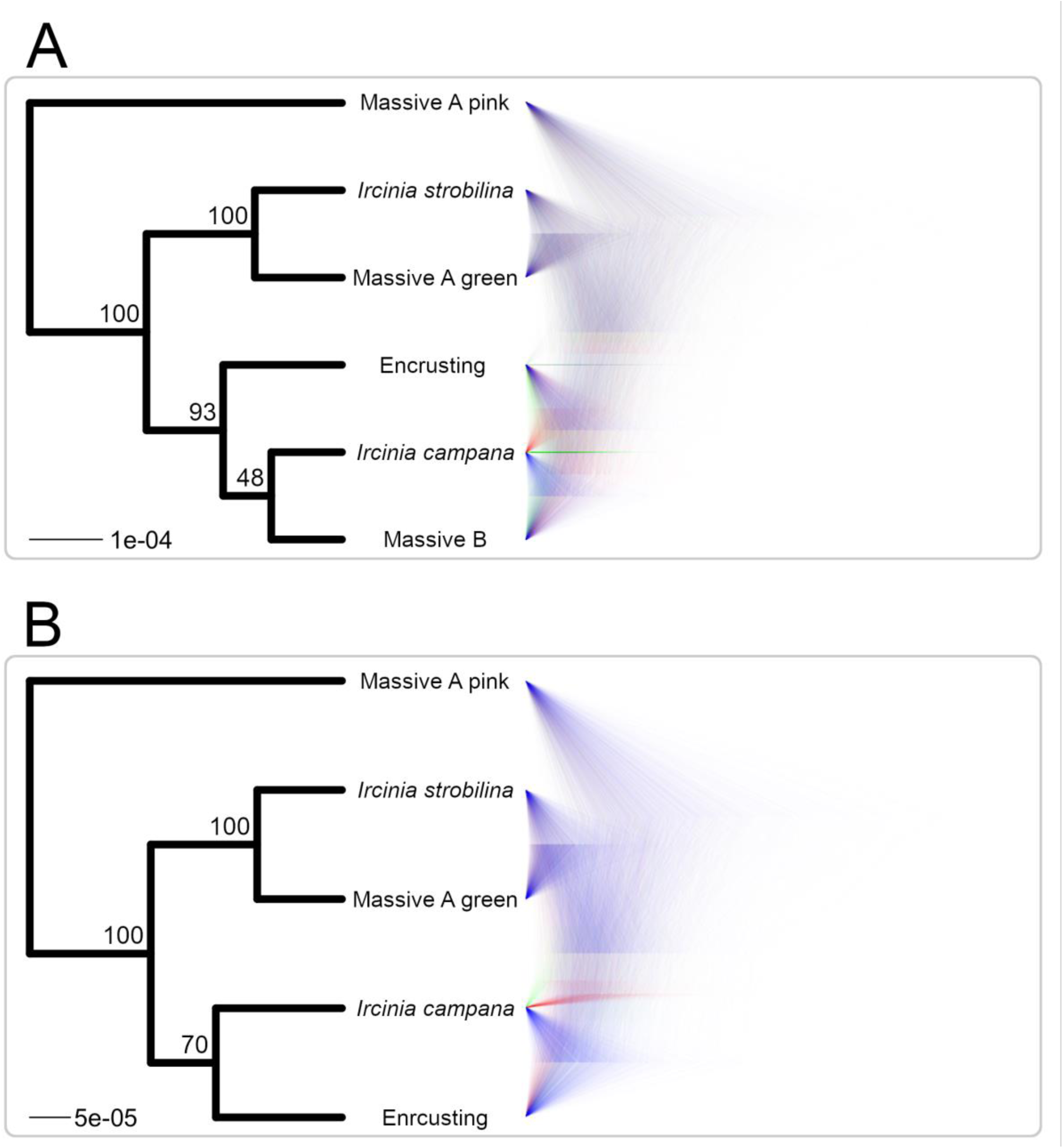
Panamanian *Ircinia* species trees inferred for the species models with the highest BFD* support using 266 2bRAD loci with SNAPP. **A** depicts species model 1 (all growth forms separate species) and **B** depicts species model 2 (split massive B by STRUCTURE populations). The left side of each subfigure portrays the maximum clade credibility consensus tree with node heights calculated as median heights, inferred in TreeAnnotator v2.5.1. Node labels are posterior probabilities. The right side shows the posterior distribution of tree as visualized in Densitree v2.6.6, in which coalescent simulations matching the dominant tree topology are drawn in blue and alternative topologies are drawn in green and red.

The estimate of ancestral populations that received the highest support by the Evanno method was K=5, followed by K =7 (Table S2, Fig. S1) (Evanno et al., 2005). The ancestry of SNPs predicted by the STRUCTURE analysis corresponded closely to source species and growth form, with the exception of the Massive B growth form (Fig. 3). However, the K=5 run identified the Massive A green and Encrusting growth forms as originating from the same ancestral population, although these two growth forms were predicted to have originated from separate ancestral populations by the K=7 run. For both the K=5 and K=7 runs, the majority of the specimens contain SNPs that were predicted to originate from an ancestral population other than the one dominating the genetic background of a given specimens’ growth form or nominal species.

**Figure 3.**
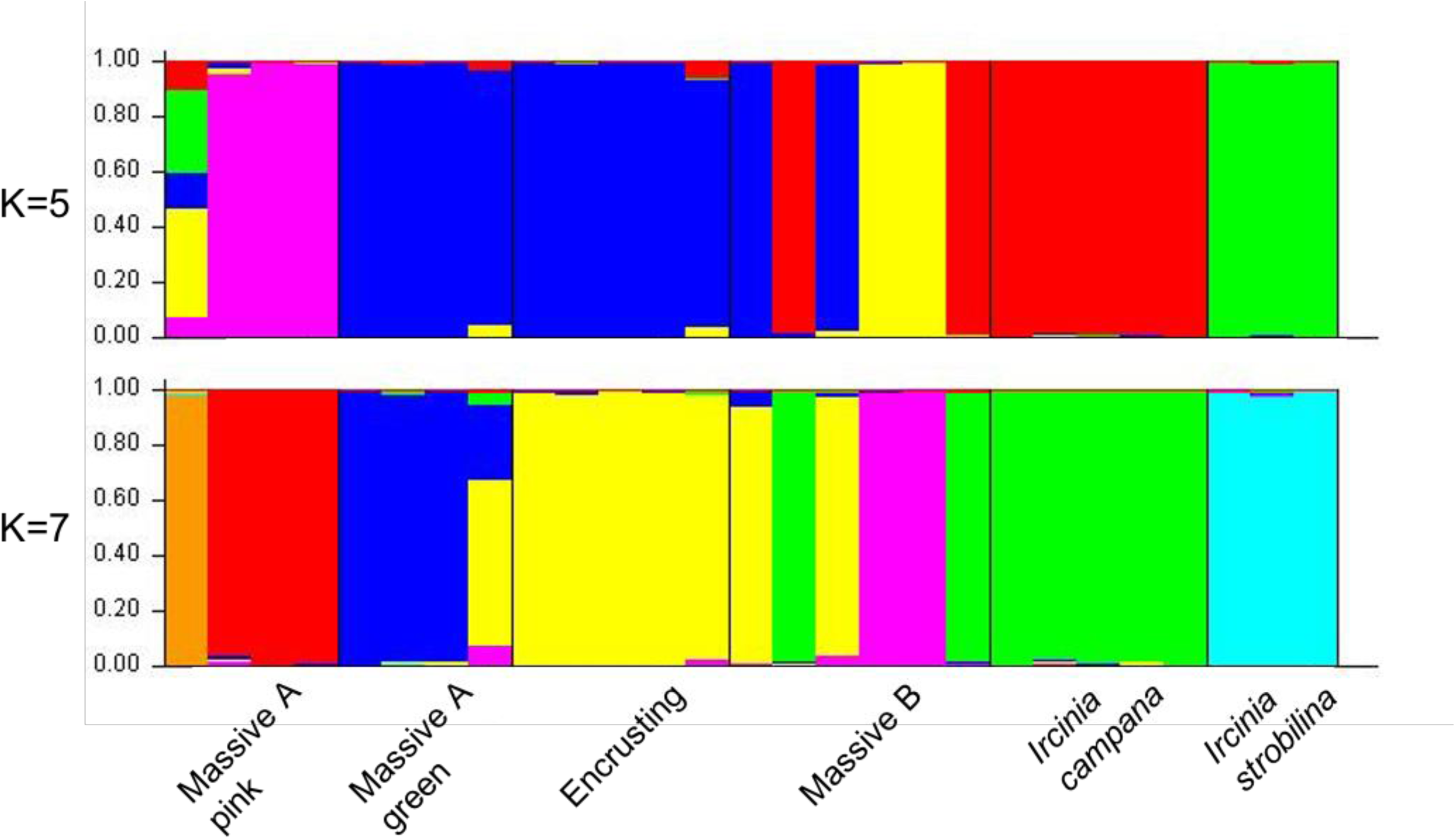
STRUCTURE plot showing assignment of SNPs to source populations, inferred for K=5 and K=7.

### Differences Among Microbiomes of the Ircinia Lineages

A total of 3,501,469 16S reads passed quality filters, including removal of chimeras, prior to clustering. 143 OTUs were identified as chloroplasts and removed from the dataset. 11938 unique prokaryotic OTUs were inferred by the mothur pipeline, with an average of 54684.38 +/- 33869.47 (1 s.d.) sequences per sample. 8612 OTUs were found only in sponges, 2612 OTUs were found only in seawater, and only 714 OTUs were shared between the two sources, a pattern consistent with observations of higher taxonomic richness in pooled sponge microbiomes relative to the environment (Thomas et al., 2016).

PERMDISP analyses on a relative abundance matrix of OTUs at the 98% clustering cutoff suggested that the assumption of homogeneity of multivariate dispersions was upheld among sponge species in model 1 (F=0.44, P=0.82) and in model 2 (F=2.22, P=0.10), although was violated when comparing seawater microbial communities and sponge microbiomes (F=43.29, P=3.90e-08). Thus, we decided to instead test for multivariate differences between sponge microbiomes and seawater microbial communities using analysis of deviance implemented in the R package mvabund, which relaxes this assumption by fitting models independently to each response variable (Y. Wang et al., 2012).

Both source (sponge vs. seawater) had a significant effect on microbiome composition (analysis of deviance: LRT = 86370.15, P = 0.001), as does host lineage (species model 1 adonis: R^2^ = 0.45, P < 1e-04; species model 2 adonis: R^2^ = 3.74, P < 1e-04). Additionally, all pairwise PERMOVAs between host taxa were significant for both species models (R^2^ values in Table S3, all P < 0.05), suggesting that each host lineage possesses microbial communities that are compositionally distinct. SIMPER analyses identified that the highest-ranking OTUs, in terms of their cumulative contributions to Bray-Curtis dissimilarities between microbiome compositions, are different for each pairwise comparison (Fig. S2). The cumulative contributions of the top 10 OTUs to Bray-Curtis dissimilarities in each pairwise comparison were substantial, and range between 22.98% (Massive A pink vs. *I. strobilina*) to 49.37% (Massive B vs. Massive A green) for species model 1 and 22.88% (Massive A pink vs. *I. strobilina*) to 44.67% (*I. campana* vs. Massive A green) for species model 2. Several of these OTUs were matched using BLASTn to ones that have been identified as being vertically transmitted in *I. felix*; however, all of these OTUs were present in the surrounding seawater, albeit in low abundances (Table S4-5, Fig. S2) (Schmitt et al., 2007). PCoAs based on Bray-Curtis dissimilarity plotted the microbiomes as clustering by host lineage in multivariate space with an average overlap of 1.21% ± 2.16% (std. dev) in SEA for species model 1 and an average overlap in SEA of 0.78% ± 2.33% (std. dev) for species model 2 (Fig. 4).

**Figure 4.**
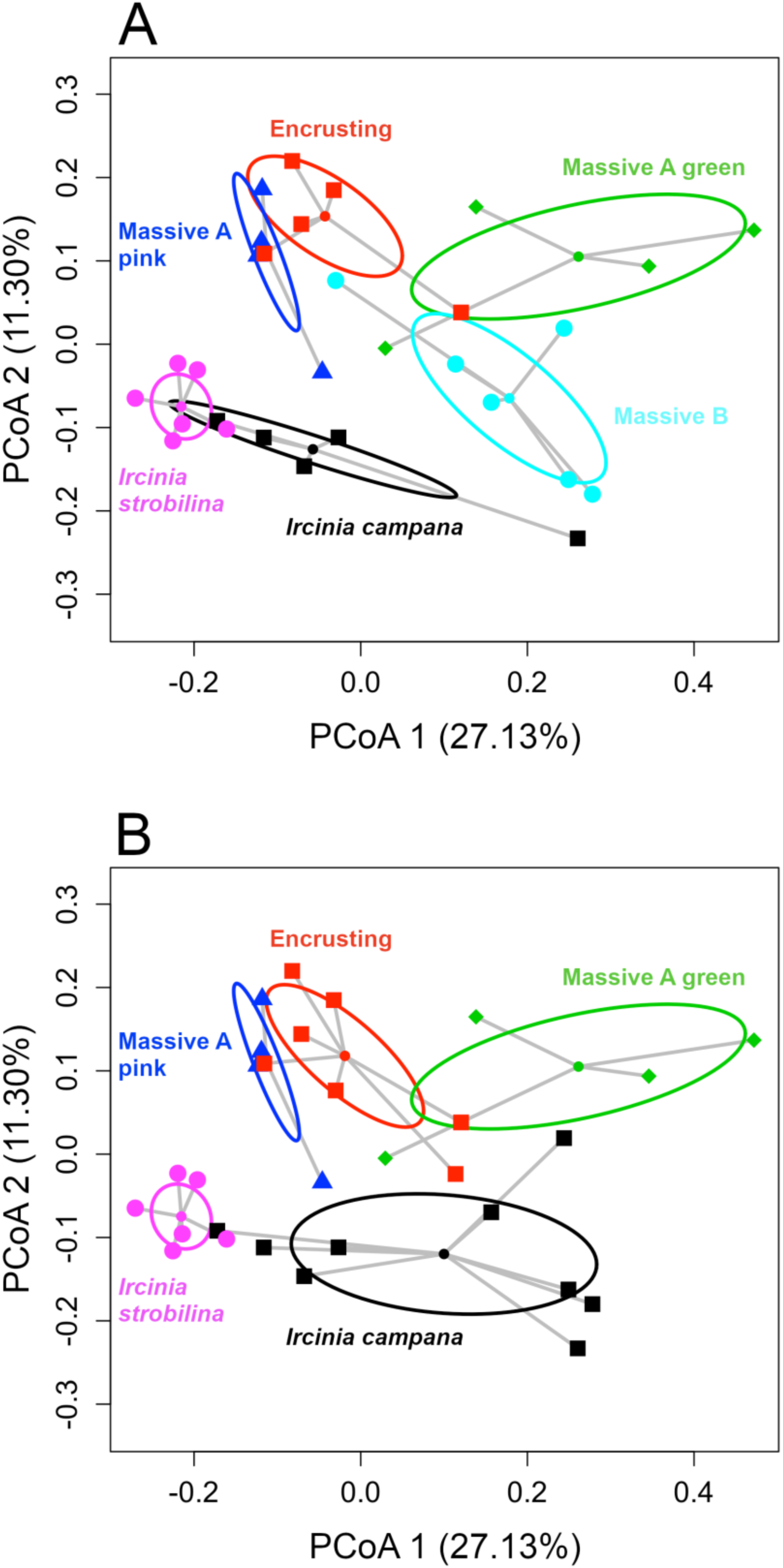
PCoAs of Panamanian *Ircinia* microbiomes constructed using Bray-Curtis dissimilarities. SEAs were calculated for host lineages inferred via BFD*, with species model 1 in **A** and species model 2 in **B**.

### ITS and CO1 Barcoding and Gene Trees

The CO1 segment amplified in 100% of the specimens and the ITS barcode amplified in roughly 90% of the specimens for which PCR was attempted. The CO1 alignment included 490 positions and the ITS alignment included 704 positions. Double peaks were found in roughly 30% of the specimens’ ITS reads, indicating possible intragenomic polymorphisms. No clear relationship between the presence of double peaks and species designation was observed. Less than 6.67 % of the pairwise CO1 comparisons and less than 3.98% of the pairwise ITS comparisons in the disparity index of pattern heterogeneity test yielded P values < 0.05 (Table S6), suggesting the sequence pairs largely evolve under the same conditions and the assumption of base composition homogeneity is upheld.

JModelTest v2.1.7 identified HKY + G as the best-fitting model of nucleotide substitution for the CO1 alignment and HKY + I + G for the ITS alignment. Chain convergence, as diagnosed by the consistent estimation of standard deviations of split frequencies below 0.01 in MrBayes v3.2.2, required 1.5 million generations per chain for both alignments. Reciprocal monophyly between individual *Ircinia* species was not observed in either gene tree (Fig. 5, 6). The CO1 sequences representing *I. campana* and the four *Ircinia* growth forms were recovered as a polytomy, which is consistent with the fact that there were no variable sites among the ingroup specimens for this gene (although 10 ‘N’ nucleotides are present in the JX306086 sequence, which is likely indicative of poor read quality). Within the ITS tree, these taxa were recovered as a poorly resolved clade with short internal branches.

**Figure 5.**
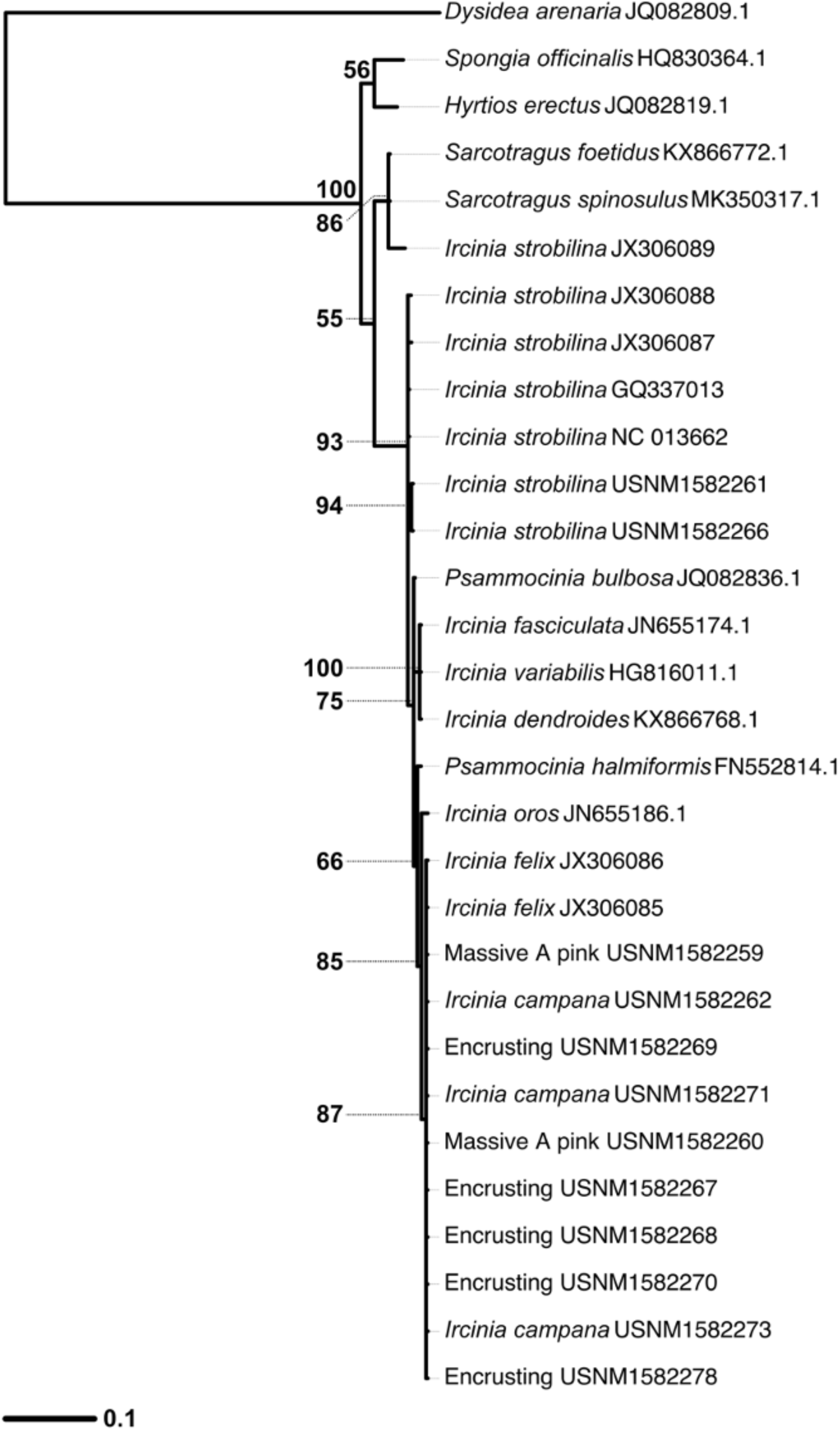
Outgroup-rooted gene tree inferred in MrBayes v3.2.2 for the CO1 alignment. The scale bar’s units are nucleotide substitutions per site, node labels are posterior probabilities, and bifurcations receiving less than 50% support are collapsed into polytomies.

**Figure 6.**
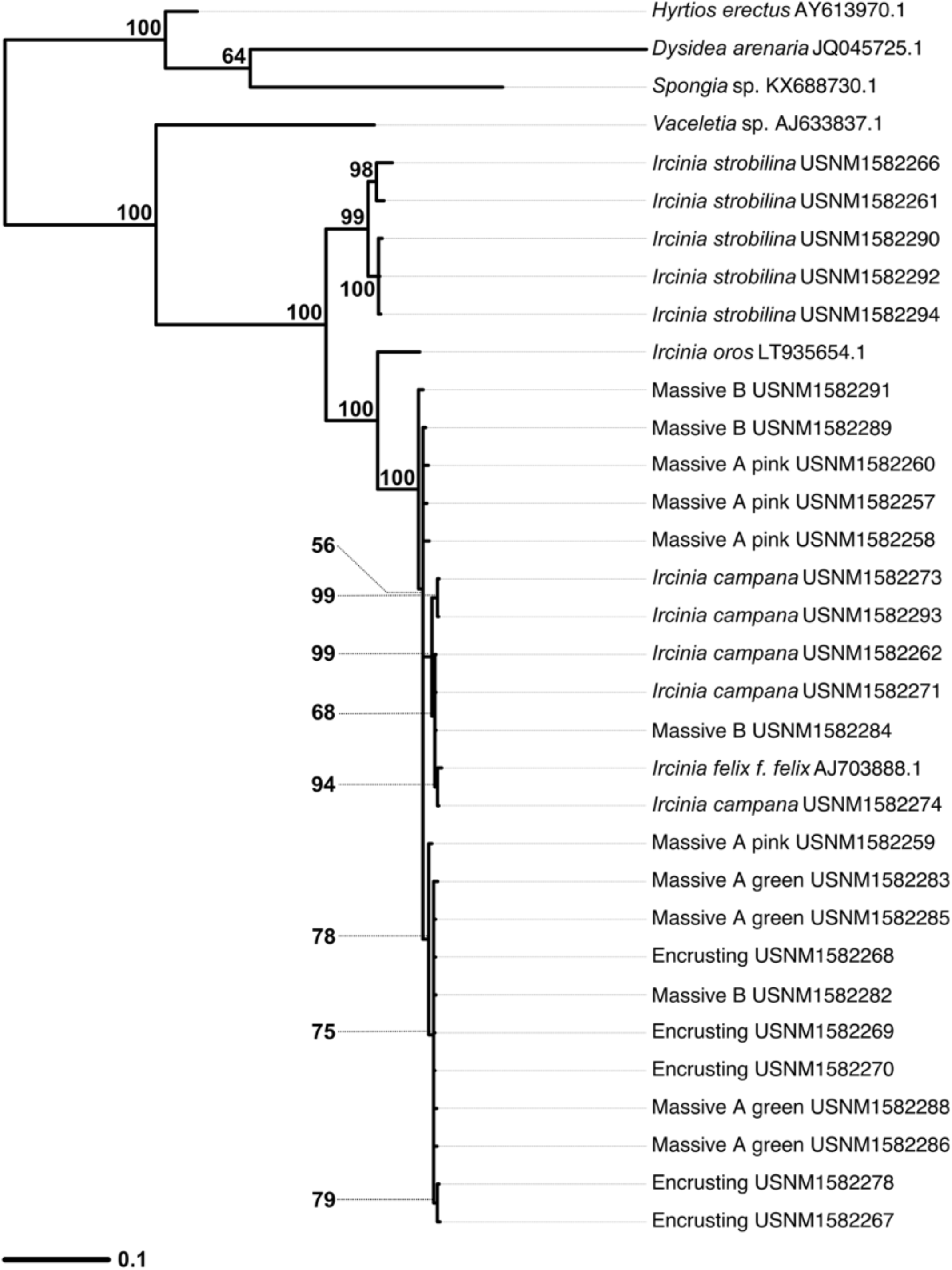
Gene tree inferred in MrBayes v3.2.2 for the ITS alignment. The scale bar’s units are nucleotide substitutions per site, node labels are posterior probabilities, and bifurcations receiving less than 50% support are collapsed into polytomies.

## Discussion

Microbiomes are recognized for their potential to unlock resource axes for their eukaryotic hosts, a role that could facilitate shifts in resource use. Here, we discovered patterns in host genetics and microbiome compositions among members of the sponge genus *Ircinia* that suggest microbiomes facilitate ecological diversification in this genus. Specifically, genetic species units that were delimited using genome-wide SNPs were found to harbor microbiomes that are compositionally distinct relative to seawater microbial communities and to the microbiomes of other *Ircinia* species with which they either share a habitat with or are in close geographic proximity to. Together, these observations suggest that the microbiomes are being actively maintained by the sponges and could be acting as agents of character displacement and local adaptation. These results provide the first test of the hypothesis that microbiomes facilitate ecological diversification among multiple, closely related (*i*.*e*. congeneric and possibly incipient) sponge species inhabiting the same locale and set the groundwork for future investigations into processes driving speciation and adaptation in sponges.

### A Comparison of Phylogenetic Analyses of the Hosts

Our analyses of genome-wide SNP data supported the hypothesis that genetic boundaries exist among the *Ircinia* growth forms and upheld species designations of the nominal species *I. strobilina* and *I. campana*. BFD* lent the highest support to two competing species models, one representing each of four growth forms and the two nominal species as distinct species (model 1), and another that differs from the first by splitting the Massive B growth form between the Encrusting growth form and *I. campana* (model 2). These results suggest that different species of *Ircinia* might be recognizable by macroscopic differences in their bodies’ growth morphologies (i.e. conule heights, oscula sizes, and pinacoderm coloration); however, caution should be exercised in the field as plasticity in some species might mistakenly lead to erroneous species identifications. *Ircinia* are well known as being taxonomically recalcitrant owing in part to variable phenotypes (de C. Cook & Bergquist, 1999), a feature that might have precipitated the incomplete resolution of the genotypes of Massive B individuals. Unfortunately, we are unable to speculate whether the growth forms in the current study correspond to the growth forms mentioned in several previous studies that describe differences in their chlorophyll-*a* content and other ecological disparities such as habitat preference (Diaz, 2005; Erwin & Thacker, 2007; Rützler et al., 2000) due to the lack of phenotypic reporting by these authors.

Even though the BFD* analysis was unable to completely resolve the genetic boundaries among the current *Ircinia*, this method outperformed the single-locus genetic barcodes CO1 and ITS. On both the CO1 and ITS gene trees, *I. campana* sequences were included within the clades containing the growth forms, whereas BFD* supported *I. campana* as being a distinct species. The failure of the *I. campana* CO1 and ITS sequences to form a reciprocally-monophyletic clade could be due to hybridization, which has also been found in Mediterranean *Ircinia*, and which appears to be present in Panamanian *Ircinia* (Riesgo et al., 2016), and incomplete lineage sorting, a likely possibility given the alternative topologies of the species tree estimates in the posterior distribution of the coalescent simulation. The presence of hybridization might also be evidence of relatively shallow barriers to reproductive isolation. Collectively, the patterns in the hosts’ genomes might imply that these taxa have speciated relatively recently. Should these *Ircinia* be young species, the passing of insufficient time since speciation events could preclude the evolution of segregating polymorphisms in the single-locus genetic barcodes among the host lineages delimited using BFD*.

### *An Appraisal of the Evidence of Microbiomes Facilitating Ecological Diversification in* Ircinia

The conservation within and the dissimilarities among the microbiome compositions of the host species recovered using BFD*, combined with their stark differences to seawater microbial communities, suggests that the *Ircinia* are actively maintaining their microbiomes. Despite the uncertain species status of Massive B, the microbiome compositions were distinct among the host lineages regardless of whether this growth form was designated as being a separate species (species model 1) or split between Encrusting and *I. campana* (species model 2). Thus, genetic distances among these *Ircinia* appear to correspond to dissimilarities among their microbiome compositions, a pattern congruent with previous studies in *I. campana* (Griffiths et al., 2019) and *C. delitrix* (Easson et al., 2020). Our data further support the developing hypothesis that divergence in microbiome compositions coincides with genetic splits among host sponge lineages, and here we demonstrate that this phenomenon is also active among multiple congeneric, possibly incipient, sponge species inhabiting the same immediate geographic vicinity.

What could be the process by which microbiomes drive adaptive diversification in *Ircinia?* We found that the OTUs that structure differences among the microbiome compositions are shared among the host lineages. Therefore, it is unlikely that these OTUs are driving the evolution of ecological novelty by virtue of introducing completely new metabolic functions to the hosts. Instead, we hypothesize that by actively controlling the relative population sizes of symbiotic microbes, the hosts are exploiting different resource axes. For example, one of the most abundant OTUs (OTU00001) regularly ranked as being among the most influential OTUs underlying Bray-Curtis distances among the host lineages. This OTU was identified as being *Candidatus* Synecoccocus spongiarum, a cyanobacterium that has received experimental support as contributing to host fitness, possibly by supplementing the host’s energy budget via photosynthates (Erwin & Thacker, 2008; C. J. Freeman & Thacker, 2011). The relative abundances of *Ca*. S. spongiarum across the host lineages appear to correspond to differences in abiotic environments associated with different habitat uses; in Massive A pink, a mangrove-dwelling sponge that grows sheltered amidst the entanglements of mangrove prop roots, *Ca*. S. spongiarum is found in trace relative abundances compared to its much higher relative abundances *I. campana*, Massive A green, Encrusting, and Massive B, all of which grow on exposed flat substrates that receive higher levels of light. Thus, the high relative abundances in this OTU in the latter taxa could be reflective of the hosts’ utilization of light via their microbes. There are many other OTUs that underly the differences among the microbiome compositions of *Ircinia* host lineages, and their contributions to host resource use could be further elucidated via metagenomic and experimental studies.

OTUs were discovered that were found in only one of the host lineages, and could thus represent microbes that introduce a novel adaptive metabolic function; however, all of these OTUS were found in very low relative abundances (results not shown). There is previous evidence of low-abundance microbes having ecological effects that are disproportionately large relative to their population sizes; in Lake Cadagno the purple sulfur bacterium *Chromatium okenii* was found to account for 70% of the total carbon uptake and 40% of the total ammonium uptake despite its relative abundance of 0.3% in the ambient microbial community (Musat et al., 2008). Similarly, a species of *Desulfosporosinus* that represented only 0.0006% of a peat bog microbial community was found to reduce sulfate at a rate of 4.0–36.8 nmol (g soil w. wt.)^−1^ day^−1^, accounting for a substantial portion of sulfate reduction pool (Pester et al., 2010). It is possible that a similar process is at play with the host lineage-specific, low abundance OTUs. However, it is difficult to reconcile the hypothesis that these microbes could be adaptive with the observation that they were found in minute relative abundances. A more credible hypothesis might state that the hosts are actively trying to prevent these microbes from establishing in large population sizes. Following this, it may be equally likely that these OTUs are pathogenic or otherwise antagonize host fitness.

Does inhabitation by symbionts cause changes in the genomes of the hosts that could result in speciation in *Ircinia*? Host immune systems are well recognized as being central to policing the crosstalk between hosts and microbes (Chaplin, 2010). Some of the mechanisms that are critical to microbiome control in vertebrates such as such as serine-threonine-directed mitogen-activated protein kinase, c-*jun* N-terminal kinase, and toll-like receptors are also present in sponges and stimulated by the pathogen-associated molecular pattern (PAMP) lipopolysaccharide (Böhm et al., 2001; Wiens et al., 2007). If microbiomes play a critical role in host fitness, then the host’s immune apparatus is likely under strong selective pressures. Given that each host lineage maintains a distinct microbiome composition, it may be the case that the immunity loci have undergone patterns of selection that differ by host lineage, thus producing different phenotypes of immune system components that could ultimately result in immune-mediated hybrid depression (Brucker & Bordenstein, 2012, 2013). Given that the active maintenance of species-specific microbiome compositions is a phylogenetically widespread phenomenon among sponges (Thomas et al., 2016), we advocate for the further investigation of immunity incompatibilities in driving sponge speciation.

### Conclusion

The body of literature concerning the implications of microbiomes for eukaryote evolution is ever advancing (Foster et al., 2017). In this study we discovered patterns in microbiome compositions and host genomes that support a model by which microbiomes facilitate ecological divergence in sponges. Addressing the question regarding whether or not microbiomes drive adaptive radiations in sponges will require a larger-scale study evaluating microbiome disparities and species richness among multiple symbiont-rich clades and clades that have relatively sparse microbiomes (Losos & Miles, 2002). Additionally, given the importance of eukaryotic immune system genes in microbial symbioses, the characterization of the patterns of selection that they undergo in sponges harboring high abundances of microbes could help elucidate the roles they play in facilitating divergent resource use via microbiomes.

## Acknowledgements

We thank Dr. Jackie L. Collier, Dr. Liliana Davalos-Alvarez, Jun Siong Low, David Carlson, Dr. Laurel Yohe, Jesse Collins, Patrick Wong, and Ben Slater for their input on the manuscript. We thank Scott Grimmell for preparing amplicons for the 16S sequencing run performed at Nova Southeastern University, Tyler Rice for his help operating a MiSeq machine, and Dr. Noah Palm for allowing use of his MiSeq. Additionally, we thank the staff of the Smithsonian Tropical Research Institute’s Bocas Research Station for their support on logistical aspects of the field work.

## Figure Captions

**Figure S1.**
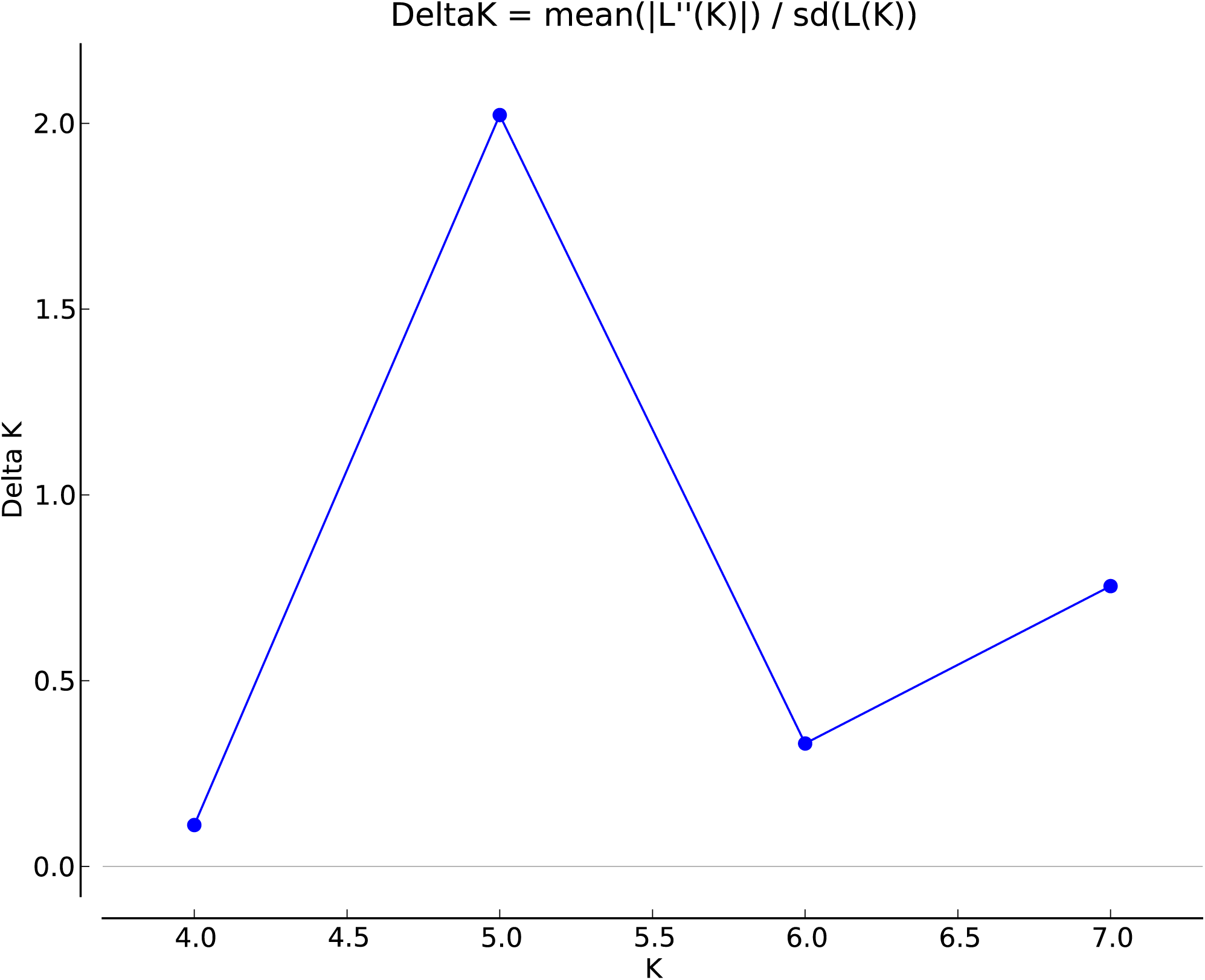
Δ*K* parameter estimates calculated using the Evanno method in Structure Harvester v0.6.94

**Figure S2.**
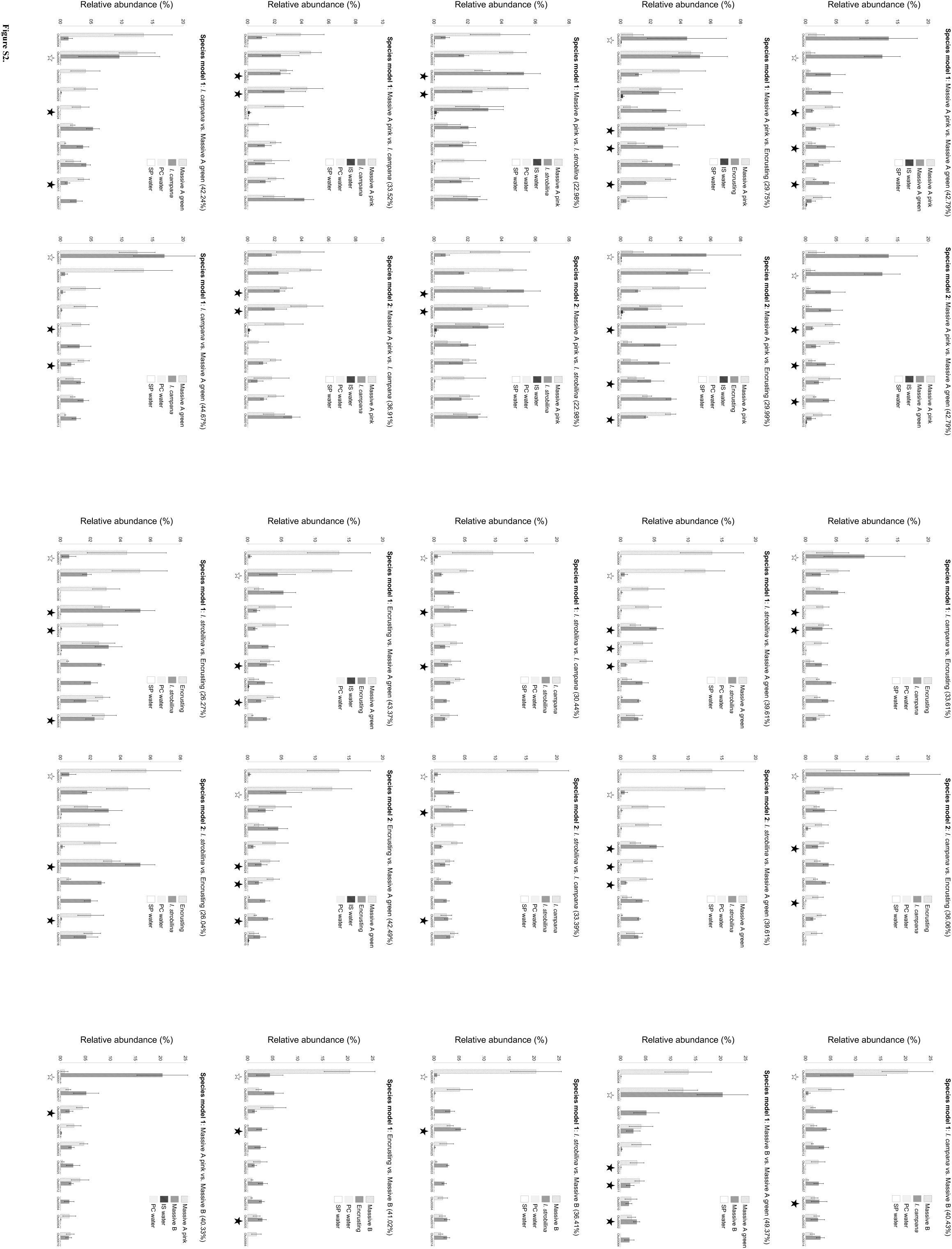
Graphs showing the relative abundances of OTUs identified by SIMPER as cumulatively contributing the most to Bray-Curtis dissimilarities. The top ten are shown for each pairwise comparison, and their cumulative contributions to Bray-Curtis dissimilarities are reported as a percent in the graph title. The relative abundances of the OTUs in the seawater samples from the sites inhabited by the sponges in each comparison are plotted alongside the sponges. Black stars denote significant BLASTn hits to vertically transmitted OTUs in Schmitt et al. 2007, and white stars denote *Candidatus* Synechococcus spongiarum. Graphs comparing the same host lineages for species model 1 and model 2 are adjacent to each other.

